# Bronchus associated lymphoid tissue induced by an attenuated *Mycobacterium tuberculosis* vaccine prevents tuberculosis from heterologous TB challenge

**DOI:** 10.64898/2026.05.21.726618

**Authors:** Dhiraj K. Singh, Garima Arora, Venkata S. R. Devireddy, Annu Devi, Mushtaq Ahmed, Vinay Shivanna, Korri Weldon, Shannan Hall-Ursone, Zhao Lai, Smriti Mehra, Xavier Alvarez, Shabaana A. Khader, Deepak Kaushal

**Author notes:** <u>To whom correspondence may be addressed:</u> Deepak Kaushal, PhD, Professor, Texas Biomedical Research Institute, 8715 W. Military Drive, San Antonio, TX 78227, USA; OR Shabaana Khader, PhD, Bernard and Betty Roizman Professor of Microbiology and Chair, Department of Microbiology, University of Chicago, Chicago, 920 E. 58th St., CLSC 1117, Chicago, IL 60637, USA. contributed equally (Dhiraj K. Singh, Garima Arora).

## Abstract

Tuberculosis (TB) remains a leading cause of death from infectious disease worldwide, underscoring the urgent need for vaccines with greater and more consistent efficacy than Bacille Calmette–Guérin (BCG). We previously showed that mucosal vaccination with an isogenic *Mycobacterium tuberculosis* (*Mtb*) mutant lacking the stress-response transcription factor SigH (*ΔsigH*) prevents pulmonary TB in two macaque species. In the absence of SigH, *Mtb* is unable to effectively counter host oxidative stress. Vaccinated macaques were notably protected from disease, exhibiting an absence of granulomatous pathology together with the formation of lymphoid follicles and robust antigen-specific CD4^+^ and CD8^+^ T cell responses, identifying *ΔsigH* as a promising live-attenuated TB vaccine candidate.

The *ΔsigH* mutant used in prior studies was generated in the *Mtb* CDC1551 background, a commonly used laboratory strain for challenge studies. Here, we evaluated whether *ΔsigH*-mediated protection extends to heterologous challenge with the more virulent *Mtb* Erdman strain. Mucosal *ΔsigH* vaccination conferred significant protection against heterologous challenge, markedly reducing pulmonary bacterial burden and TB-associated pathology. Longitudinal high-resolution PET/CT imaging demonstrated that aerosol *ΔsigH* vaccination induced robust inducible bronchus-associated lymphoid tissue (iBALT) responses in the lung. Unlike granulomas, these iBALT structures resolved over time while remaining associated with protection against subsequent *Mtb* challenge. Protection of highly susceptible rhesus macaques against virulent heterologous *Mtb* challenge following aerosol *ΔsigH* vaccination supports the further preclinical development of *ΔsigH*-based live-attenuated TB vaccines and highlights iBALT induction as a potential correlate and mechanistic driver of protective immunity against TB.

## Introduction

While impressive gains have been made in the global control of HIV over the past two decades^1^, tuberculosis (TB) remains a major global infectious disease worldwide, causing more than 1.3 million deaths annually^2^. The effectiveness of the currently licensed vaccine for TB prevention, Bacille Calmette-Guerin (BCG), in preventing adult pulmonary TB, is limited^3^, highlighting the urgent need for improved vaccination strategies. Effective TB vaccines must elicit robust and durable lung T cell immunity capable of controlling *Mycobacterium tuberculosis* (*Mtb*) infection^4^.

Live-attenuated *Mtb* strains induce a broader repertoire of antigenic responses compared with BCG^5,6^. Despite this promise, only one live-attenuated *Mtb* vaccine candidate, MTBVAC (*ΔphoP*/*ΔfadD26*), has advanced to human clinical testing till now^7,8^. Attenuated *Mtb* vaccines lacking key virulence or detoxification factors not only potentially develop durable immunity but offer a system where the replication of the live vaccine strain is rapidly restricted, often by innate immunity, prior to the development of granulomatous pathology. The alternative sigma factor SigH plays a central role in protecting *Mtb* from multiple host-induced stresses, including oxidative stress^9,10,11,12,13,14,15,16,17^. Lack of SigH causes a markedly reduced survival and pathogenicity in the lungs^18^, even under conditions of immunodeficiency^19^. These findings demonstrate that SigH-regulated transcriptional program is critical for *Mtb* persistence within the host lung. Consistent with this, transcriptomic analyses of bacilli recovered from infected phagocytes show increased expression of *sigH*^15^. Further supporting its role in virulence, *Mtb* strains containing genomic duplications encompassing the *sigH* locus exhibit enhanced pathogenicity in the lungs^20^. Similarly, increased *sigH* copy number has been shown to reduce the protective efficacy of BCG in animal models^21^. Together, these observations suggest that loss of SigH function is strongly associated with the induction of protective host immune responses. Consistent with this hypothesis, macrophages infected with *ΔsigH* exhibit increased apoptosis and autophagy compared with those infected with wild-type *Mtb*^22^.

Aerosol infection of rhesus macaques with *ΔsigH* induces robust innate and adaptive immune responses that confer protection against subsequent *Mtb* challenge^23^. *ΔsigH* infection is associated with enhanced antigen presentation and the recruitment of Type I IFN–conditioned T cells, which display elevated Th1 and Th17 responses compared with BCG vaccination, ultimately promoting more efficient macrophage-mediated killing of *Mtb*^24^ and signatures of B-T cell cooperation^24^. The current study had two objectives: I) Test if aerosol vaccination with *ΔsigH* can protect susceptible rhesus macaques against challenge with a heterologous strain of *Mtb* (i.e., a strain different from the one in which the *sigH* deletion was generated). We have previously shown that compared with CDC1551, *Mtb* Erdman exhibits substantially greater pathogenicity in rhesus macaques, characterized by significantly higher bacillary replication in lungs and granulomas as well as dysregulated recruitment of myeloid cells^25^; and II) Employ live imaging to study if iBALT is recruited post-*ΔsigH*-vaccination and to characterize these early iBALT responses vis-a-via TB granulomas in detail and to understand the mechanisms operation in iBALT which generate protective responses.

## Methods

### NHP study design and infections

20 mycobacteria-naive rhesus macaques (RMs)^26^ sourced from the Southwest National Primate Research Center, were used (**Table S1**). All procedures adhered to NIH guidelines and received approval from the Institutional Animal Care and Use Committees (IACUC) of Texas Biomedical Research Institute. Specifically, the animals were either unvaccinated (n=7) vaccinated intradermally with BCG (n=7) or aerosol vaccinated with *ΔsigH* (n=6), as described earlier^23^. 12-weeks post-vaccination, macaques were exposed to 100 CFU aerosolized *Mtb* CDC1551 (**Fig 1a**). Infection was assessed through tuberculin skin test (TST) or antigen-specific ICS, while TB progression was monitored by longitudinal measurements of weight, temperature, and C-reactive protein (CRP) and bronchoalveolar lavage (BAL) CFUs, and PET/CT scans as described^23,27-32^. Dissemination was evaluated during necropsy by culturing bronchial lymph node, spleen, liver, and kidney tissues to measure CFUs. Demographic information including age, gender, etc., and study specific information of macaques are provided (**Table S1**). Animals were euthanized at 12-13 weeks post-challenge.

### Sampling

TST was performed 1-3 weeks before challenge and at weeks 3 and 5 post-challenge, as described^23,33^. PET/CT scans were performed at week 4 post-challenge and at the endpoint as described earlier and compared to baseline^30-32^. BAL samples were obtained one week before either vaccination or *Mtb* infection and subsequently every two weeks, as described^23,34^. BAL cells were used for determining bacterial burden and cellular analysis through flow cytometry, as described^23,34^. Blood samples were collected one week prior to vaccination or *Mtb* infection and thereafter on a weekly basis, for measuring complete blood count, serum chemistry, including serum C-reactive protein (CRP), and for flow cytometry, employing the flow panels specified earlier^29,34,35^.

**Figure 1.**
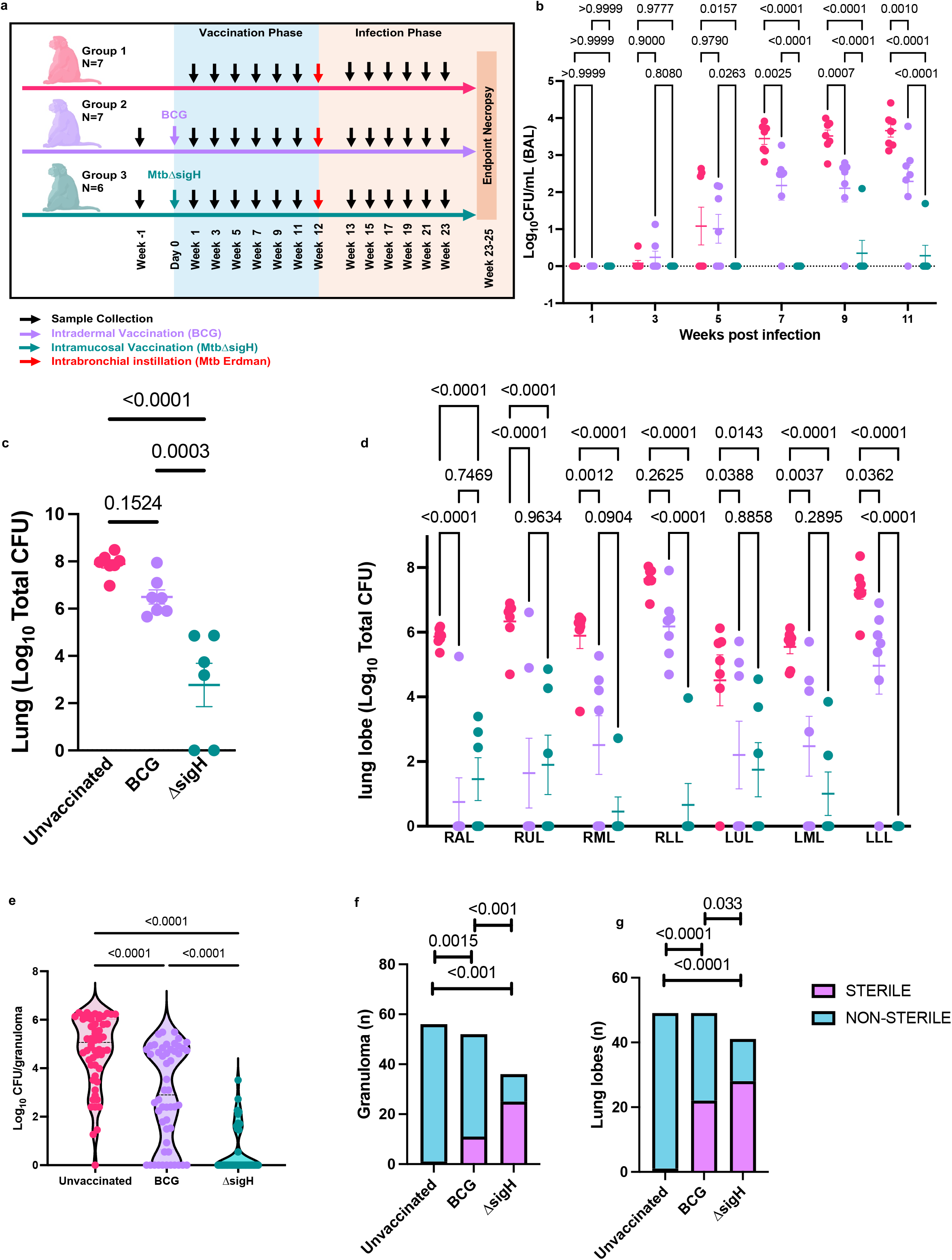
Measurement of TB. (**a**) Experimental design. In unvaccinated (strawberry), BCG (lavender) and *ΔsigH*-vaccinated (teal) CMs post-*Mtb* infection, shown are endpoint serum Albumin/Globulin (A/G) ratios (**b**); blood Neutrophil (N)/Lymphocyte (L) ratios (**c**) and CXR scores (**d**); longitudinal BAL Colony Forming Units (CFUs) (**e**); endpoint lung (**f**), lung granuloma (**g**), and bronchial lymph node (**h**) CFUs per gram. Gross pathology in representative unvaccinated (**i**), BCG- (**j**) and *ΔsigH*-vaccinated (k) with granulomas (black arrows). Representative sub-gross histopathology and histopathology of unvaccinated (**l, o**), BCG- (**m, p**) and *ΔsigH*-vaccinated (**n, q**) CM lungs. Unvaccinated: Multiple confluent necrotic granulomas (**o**), BCG-vaccinated: (**p**) and *ΔsigH* -vaccinated: Solitary non-necrotic small granuloma with lymphoid hyperplasia and iBALT (**q**). Morphometric lung pathology score in the three groups (**r**). Percentage of cellular inflammation (**s**), necrotic region (**t**) and lymphoid aggregates (**u**), across multiple fields of view of lung tissue using HALO (Indica Labs). Data is presented as mean ± standard error of mean (SEM) and P-values were calculated by one-way ANOVA with Tukey’s correction, except for (**d**) where a two-way ANOVA with Tukey’s correction was used.

### Tissue bacterial burden and pathology

Tissues were collected and processed as described^23^. CFUs were determined per gram of tissue and per mL of BAL fluid. Lung pathology at necropsy was assessed by a board-certified veterinary pathologist in a blinded manner, utilizing zinc-formalin-fixed paraffin-embedded (FFPE) tissues representing all lung lobes using previously described methods^23^.

## Results

### Post-vaccination safety and clinical parameters

We performed an in-depth safety analysis in 20 rhesus macaques unvaccinated controls, n = 7; BCG-vaccinated [1×10^6^ CFU, intradermal], n = 7; *ΔsigH*-vaccinated [1×10^3^ CFU, aerosol], n = 6) (**Fig 1a**). All animals were tuberculin skin test (TST)-negative prior to vaccination and converted to positive five weeks post-*Mtb* challenge. No TST was administered during the vaccination phase (**Table S1**). Importantly, none of the animals displayed any clinical signs of active infection or disease following vaccination, including dyspnea, anorexia, pyrexia (**not shown**) or loss of body weight (**Fig S1a**). In addition, no significant changes were observed in the acute phase marker serum C-reactive protein (CRP) (**Fig S1b**), and the peripheral blood indicators associated with TB disease progression in macaques^36^, including serum A/G (**Fig S1c**) or peripheral blood neutrophil/lymphocyte (N/L) ratio (**Fig S1d**) or frequency of blood neutrophil (**Fig S1e**) and lymphocyte levels (**Fig S1f**), during the *ΔsigH* vaccination phase relative to BCG vaccination.

Furthermore, PET/CT scans performed six and eleven weeks after vaccination for all 6 *ΔsigH* vaccinated animals showed no granulomatous abnormalities (**Fig S1g**). At weeks 7, 9 and 11, post-vaccination, we studied *ΔsigH*-bacillary burdens in bronchoalveolar lavage (BAL) samples. Only one out of six *ΔsigH*-vaccinated macaque showed viable bacilli at one time-point (week 9) (**Fig S1h**), albeit at very low levels (∼2 logs); whereas no *ΔsigH* bacilli were detectable at weeks 7 or 11 post-vaccination in any animals or in five of six animals at week 9 (**Fig. S1h**).

### Significant reduction of TB in rhesus macaques vaccinated with *ΔsigH* via the mucosal route following heterologous *Mtb* Erdman challenge

Twenty rhesus macaques in the unvaccinated, BCG-vaccinated and *ΔsigH*-vaccinated groups, all underwent challenge with ∼20 CFU of *Mtb* Erdman via intra-bronchial instillation, 12 weeks post-vaccination (**Fig 1a**). To assess the impact of impact of *ΔsigH* vaccination on heterologous *Mtb* infection, bacillary replication in the lungs was first monitored using was first measured on BAL as a surrogate, analyzed by two-way ANOVA with Tukey’s multiple comparison correction.

Significantly lower *Mtb* burdens were recovered from BAL samples of *ΔsigH*-vaccinated macaques compared with both unvaccinated (*P*=0.0157, wk 5; *P*<0.0001, wk 7; *P*<0.0001, wk 9; *P*<0.0001, wk 11) and BCG-vaccinated animals (*P*=0.0263, wk 5; *P*<0.0001, wk 7; *P*<0.0001, wk 9; *P*<0.0001, wk 11) beginning week 5 post-challenge and continuing through the end of the study (**Fig 1b**). BCG vaccination also resulted in significantly reduced BAL *Mtb* burdens relative to unvaccinated controls at weeks 7–11 (P = 0.0025, week 7; P = 0.0007, week 9; P = 0.0010, week 11) (**Fig 1b**). The lack of significant differences at earlier time-points likely reflects the low initial bacillary burdens associated with to low-dose intrabronchial *Mtb* challenge, in contrast to the higher inoculum used in previous aerosol challenge studies^23,24^.

Consistent with these findings, *ΔsigH* vaccination markedly reduced bacterial burdens throughout the respiratory tract. Significantly lower *Mtb* CFU were recovered from whole lungs (**Fig 1c**), individual lung lobes (**Fig 1d**), lung-derived granulomas (**Fig 1e**), lung-draining bronchial lymph nodes (BrLN) (**Fig S2a**), axillary lymph nodes (AxLN), spleen, liver and kidneys (**Fig S2b-e**) of *ΔsigH*-vaccinated macaques, relative to both the other groups. The mean lung *Mtb* burden in *ΔsigH-*vaccinated macaques was ∼5.1-logs lower than in unvaccinated (animals *P*<0.0001), and ∼3.7-logs lower than in BCG-vaccinated animals (*P*<0.0003) (**Fig 1c**). While intradermal BCG vaccination resulted in ∼1.4 log lower CFU than unvaccinated animals, this reduction did not reach statistical significance relative to unvaccinated animals (*P*=0.1524) (**Fig 1c**). Taken together, these results are comparable to our previous observations in rhesus^23^ and cynomolgus macaques^24^ where the efficacy of aerosolized *ΔsigH* vaccination was assessed following homologous *<u>Mtb</u>* CDC1551 challenge. Because the *Mtb* challenge here was delivered into the right lower lung lobe, this and the contralateral left lower lobe harbored the highest bacillary burdens in unvaccinated animals (6.15–6.3 log CFU) (**Fig 1d**). In both lobes, bacterial burdens were significantly reduced in *ΔsigH*-vaccinated animals compared with both unvaccinated and BCG-vaccinated groups (P < 0.0001 in all comparisons). Bacillary burdens in the remaining lobes were lower in unvaccinated animals (3.9–5.8 log CFU) compared to those in the right and the left lower lobes (**Fig 1d**). Both BCG and *ΔsigH* vaccination significantly reduced bacterial loads in these lung lobes. We next quantified *Mtb* CFU in individual granulomas (**Fig 1e**). Granulomas from unvaccinated animals harbored a mean burden of 4.71 logs, whereas those from the BCG- and the *ΔsigH*-vaccinated macaques contained significantly lower burdens of 2.87 log- and 0.59 log-CFU, respectively (*P*<0.0001 in both cases). Importantly, granulomas from *ΔsigH*-vaccinated animals also contained significantly fewer bacilli than those from BCG-vaccinated animals (P < 0.0001). We further evaluated the frequency of sterile lesions defined as either lung lobes (**Fig 1f**), or individual granulomas (**Fig 1g**) from which culturable *Mtb* could not be recovered. While almost all lung lobes from unvaccinated macaques contained culturable *Mtb* (98%), this proportion decreased to 55 and 31%, respectively, for BCG- and *ΔsigH*-vaccinated macaques (**Fig 1f**). Similarly, none of the granulomas from unvaccinated macaques were sterile, whereas approximately 20% of granulomas from BCG-vaccinated animals and ∼70% from *ΔsigH*-vaccinated animals were sterile (**Fig 1g**). These differences were highly significant by Fisher’s exact test (P = 0.0015 for unvaccinated vs. BCG; P < 0.001 for unvaccinated vs. *ΔsigH*; P < 0.001 for BCG vs. *ΔsigH*). Consistent with previous reports showing that *Mtb* burdens can accumulate to even higher levels in bronchial lymph nodes (BrLN) than in lungs on a per-gram basis, we observed substantial bacterial burdens in BrLN of unvaccinated animals. In contrast, BrLN from *ΔsigH*-vaccinated macaques were sterile, with bacterial burdens reduced by approximately 5 logs relative to unvaccinated animals and ∼4.5 logs relative to BCG-vaccinated animals (**Fig S2a**). Protection conferred by *ΔsigH* vaccination also extended to extrapulmonary tissues, including axillary lymph nodes (AxLN), spleen, liver, and kidneys (**Fig S2b-e**). These tissues were largely sterile in *ΔsigH*-vaccinated animals, whereas substantial bacterial burdens were present in unvaccinated and BCG-vaccinated animals. For example, mean AxLN burdens were 2.43, 1.1, and 0 log CFU in unvaccinated, BCG-vaccinated, and *ΔsigH*-vaccinated animals, respectively, with the difference between unvaccinated and *ΔsigH* groups reaching statistical significance (P = 0.0010). Similar patterns were observed in spleen, liver, and kidney tissues, with significantly reduced bacterial burdens in *ΔsigH*-vaccinated animals relative to controls. Collectively, these microbiological findings demonstrate profound suppression of *Mtb* replication across pulmonary and extrapulmonary compartments following *ΔsigH* vaccination, even after challenge with the highly pathogenic *Mtb* Erdman strain.

### Clinical indicators corroborate protection following *ΔsigH* vaccination

The profound reduction in bacterial replication in *ΔsigH*-vaccinated macaques was mirrored by clinical parameters associated with progressive TB disease post-*Mtb* challenge. By the end of the study, *ΔsigH*-vaccinated macaques exhibited a mean body weight gain of approximately 10%, whereas BCG-vaccinated animals showed no significant weight gain and unvaccinated animals lost approximately 5% of body weight (**Fig 2a**). These differences were statistically significant. Peripheral blood immune profiles also differed significantly between groups. Lymphocyte frequencies were significantly higher in *ΔsigH*-vaccinated macaques at the endpoint post-challenge compared with BCG-vaccinated and unvaccinated animals (P = 0.0174 and P = 0.0042, respectively) (**Fig 2b**). Conversely, neutrophil frequencies were significantly lower in *ΔsigH*-vaccinated animals at the endpoint relative to both comparison groups (P = 0.0186 and P = 0.0072, respectively) (**Fig 2c**). Together these resulted in lower N/L ratios in *ΔsigH*-vaccinated, relative to BCG-vaccinated or unvaccinated macaques at the endpoint (**Fig S2f**). These parameters were also significantly different between the unvaccinated/BCG-vaccinated and *ΔsigH*-vaccinated groups at several other timepoints post-*Mtb* infection (**Fig S2g-i**). Markers of systemic inflammation further supported these findings. Albumin-to-globulin (A/G) ratio, which decreases during progressive TB disease^36^, was significantly reduced in unvaccinated and BCG-vaccinated animals compared with *ΔsigH*-vaccinated macaques (P = 0.0001 and P = 0.0005, respectively) at the endpoint (**Fig 2d**), due to both a significant decline in serum albumin levels (**Fig 2e**) and a significant increase in serum globulin levels (**Fig 2f**) in unvaccinated or NCG-vaccinated macaques, relative to those vaccinated with *ΔsigH*. These parameters were significantly different between unvaccinated/BCG-vaccinated and *ΔsigH*-vaccinated groups not only at the endpoint but at several other timepoints post-*Mtb* challenge (**Figs S2j-l**). Whereas Serum C-reactive protein (CRP) levels were elevated in unvaccinated and BCG-vaccinated macaques relative to *ΔsigH*-vaccinated animals at the study endpoint (**Fig S2m**), these levels were significantly reduced in *ΔsigH*-vaccinated animals at week 11 post-challenge (**Fig 2g**). Together, these clinical indicators corroborate the microbiological findings and demonstrate that *ΔsigH* vaccination confers strong protection against heterologous challenge with the highly pathogenic *Mtb* Erdman strain in rhesus macaques.

**Figure 2.**
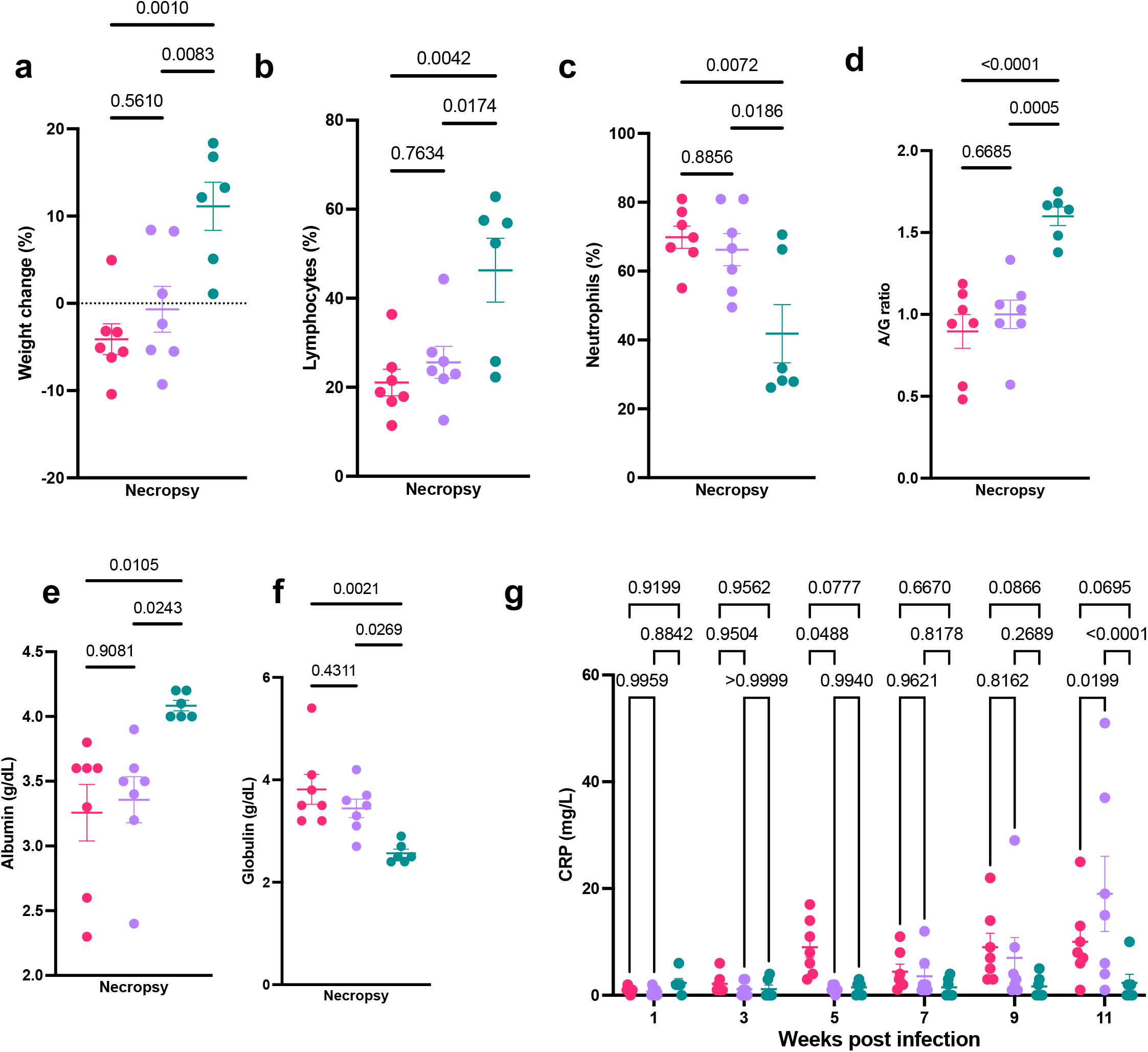
Lymphocytic immune responses in BAL post-vaccination. Absolute counts of CD3^+^ (**a**), CD4^+^ (**b**), and CD8^+^ (**c**) T cells and B cells (**d**), in BAL at weeks 3, 5, and 7b post-vaccination time-point in BCG- (lavender) or *ΔsigH-* (teal) vaccinated CMs. Each column represents an individual macaque (n=5). Frequencies of CCR5^+^ and CXCR3^+^ - CD3^+^ (**e, I**), CD4^+^ (**f, j**), and CD8^+^ (**g, k**) T cells and B cells (**h, l**) in BAL. Frequencies of CCR6^+^ CD3^+^ (l), CD4^+^ (**m**) and CD8^+^ (**n**) T cells in BAL. Frequencies of CXCR3^+^CCR6^+^ CD4^+^ (**o**) and CD8^+^ T cells in BAL (**p**); effector (**q**), memory (**r**) and naïve (**s**) CD4^+^ T cells, CD69^+^ (**t**) and KI67^+^ (**u**) effector CD4^+^ T cells, CCR5^+^ (**v**), CCR7^+^ (**x**) and KI67^+^(**z**) memory CD4^+^ T cells and CCR5^+^ (w), CCR7^+^ (**y**) and KI67^+^(**a’**) memory CD8^+^ T cells in BAL at weeks 3, 5, and 7 post-vaccination time-point, expressed as percentage of parental population. P-values are derived from multiple Mann-Whitney tests with multiple hypothesis correction by false discovery rate method of Benjamini, Kreiger and Yekutielli (two stage step up). Results are shown for weeks 3, 5, and 7 post-vaccination time-point (n=5). Each column represents an individual macaque.

### *ΔsigH* vaccination reduces lung pathology following *Mtb* Erdman challenge

Gross pathological examination at necropsy revealed extensive granulomatous and inflammatory involvement in the lungs of unvaccinated macaques (**Fig 3a,d**), whereas BCG-vaccinated (**Fig 3b,e**) and *ΔsigH*-vaccinated animals (**Fig 3c,f**) exhibited markedly reduced pathological changes. Sub-gross histopathological analysis confirmed these observations. Both *ΔsigH*-(**Fig 3i, l**) and BCG-(**Fig 3h, k**) vaccinated animals displayed significantly fewer lung lesions, along with reduced edema, pneumonia, and diffuse inflammatory infiltrates following *Mtb* challenge compared with unvaccinated macaques (**Fig 3g–j**). Both vaccinated groups had significantly lower total morphometric quantitative lung pathology scores (**Fig 3m**) and extent of necrosis (**Fig 3n**) than unvaccinated animals (P < 0.0001), although no statistically significant difference could be detected between the BCG- and *ΔsigH*-vaccinated groups. in terms of necrosis. Lesions in unvaccinated animals were frequently large, confluent, and centrally necrotic, with occasional smaller non-necrotic lesions adjacent to larger necrotic granulomas. In contrast, vaccinated macaques retained a greater proportion of normal lung architecture and developed smaller granulomas. BCG-vaccinated animals more frequently exhibited non-necrotic granulomas compared with unvaccinated controls. Strikingly, all but one *ΔsigH*-vaccinated macaque lacked detectable granulomas in the lungs. In the few lesions observed in *ΔsigH*-vaccinated animals, granulomas were non-necrotic and enriched in lymphoid follicles, consistent with the enhanced formation of iBALT structures. The lungs of *ΔsigH*-vaccinated animals had significantly greater lymphocytic aggregates or follicles than each of the other two (unvaccinated, BCG-vaccinated) groups (**Fig 3o**) and this group also exhibited greater cellular inflammation than the BCG group (**Fig 3p**), although the inflammation was significantly lower than in the unvaccinated group. We have previously shown that the formation of lymphoid follicles into inducible bronchus associated lymphoid tissue, is a hallmark of *ΔsigH*-mediated protection against TB. Collectively, these findings demonstrate that *ΔsigH* vaccination markedly reduces lung pathology following heterologous *Mtb* Erdman challenge and promotes the formation of organized lymphoid structures associated with protective immunity.

**Figure 3.**
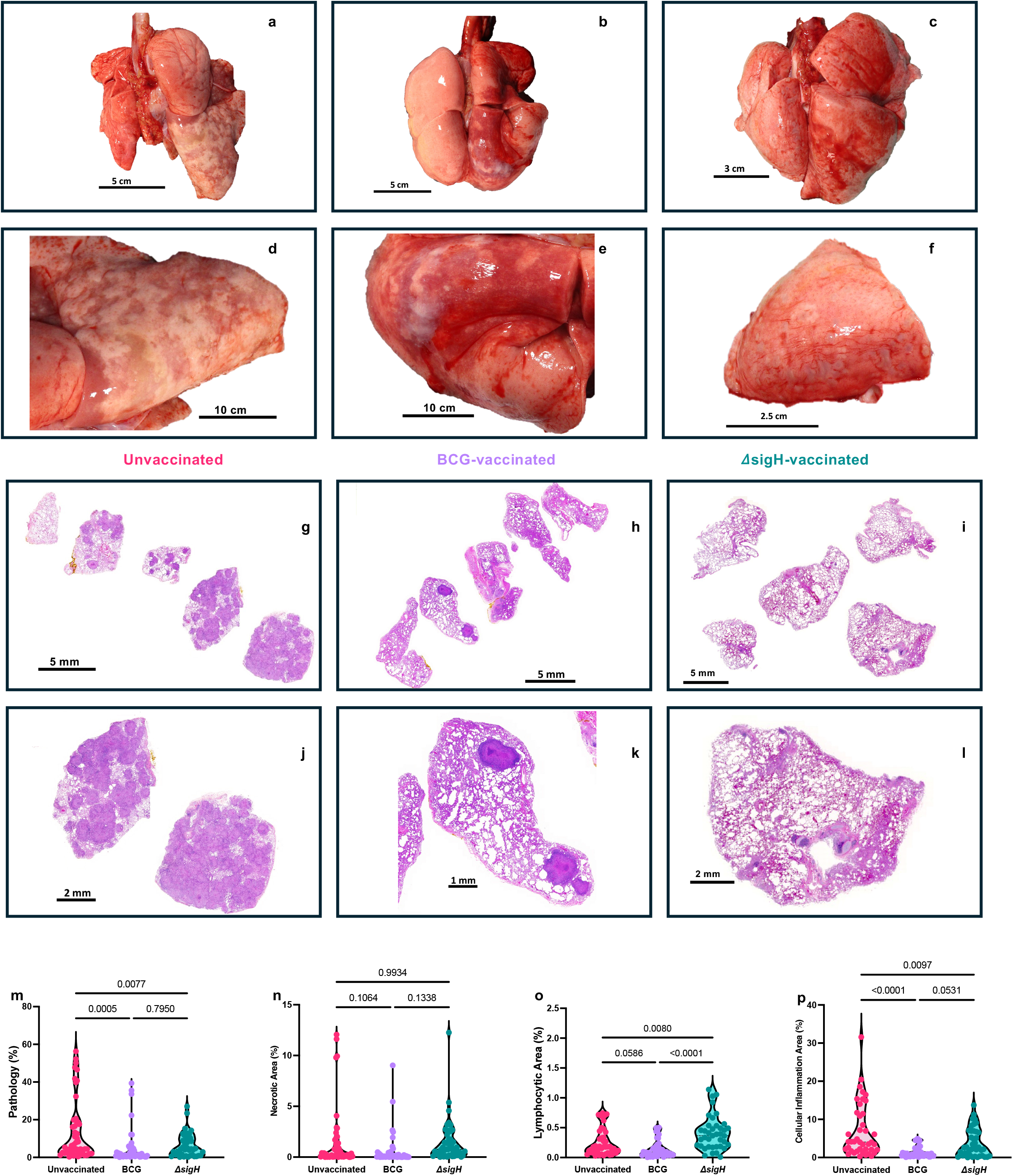
Antigen-specific ICS BAL post-vaccination but pre-challenge. Frequencies of CD4^+^ and CD8^+^ T cells expressing IFNG (**a, f**), TNF-α (**b, g**), IFNG and TNF-α (**c, h**), GZMB (**d, i**) and IL17 (**e, j**) in response to *Mtb* Cell-Wall (CW) fraction in CMs vaccinated with BCG (lavender) or *ΔsigH* (teal). Frequencies of CD4^+^ and CD8^+^ T cells expressing IFNG (**k, p**), TNF-α (**l, q**), IFNG and TNF-α (**m, r**), GZMB (**n, s**) and IL17 (**o, t**) in response to pooled peptide pools of *Mtb* ESAT6 and CFP10 (EC). Each column represents an individual macaque (n=5) at weeks 3, 5, and 7 post-vaccination. P-values are derived from multiple Mann-Whitney tests with multiple hypothesis correction by false discovery rate method of Benjamini, Kreiger and Yekutielli (two stage step up).

We also measured the extent of lung pathology by high-resolution PET/CT scans of these groups of macaques after injecting with 18FDG radioisotope (**Fig 4**). PET/CT was performed at two timepoints post-*Mtb* infection (weeks 4-6 and weeks 12-13, the latter being just prior to the euthanasia for each animal). Rendering of granulomatous pathology observed in PET/CT scans clearly indicates the formation of numerous granulomas in unvaccinated animals at week 4-6 (**Fig 4a**), which occupied considerable lung volume, despite animal-to-animal heterogeneity. Both the number and the volume of granulomas was reduced by BCG vaccination at week 4 (**Fig 4b**), and even further reduced by *ΔsigH* vaccination, at week 4-6 (**Fig 4c**). In our renderings, bronchial lymph nodes are colored in red, and most signal in the lungs of *ΔsigH* vaccinated macaques at 4-6 weeks was localized in this tissue. Quantitative analysis of SUV^max^ (maximal standardized uptake value) suggested significant reduction after 4-6 weeks of *Mtb* Erdman challenge in both vaccination groups relative to unvaccinated macaques, but there were no statistically significant differences between the BCG- and the *ΔsigH*-vaccinated groups (**Fig 4d**). The number of lesions detected by PET/CT and volume occupied by them was also significantly lower at week 4-6 in both vaccination groups relative to unvaccinated macaques (**Fig S3a, b**). At weeks 12-13, i.e., just prior to end of protocol, the lungs of all six unvaccinated macaques exhibited massive granulomatous pathology by PET/CT scans and demonstrated a massive increase in both the number and the volume occupied by granulomas relative to week 4-6 (**Fig 4e**). Substantially increased granulomatous pathology over time was also apparent in the BCG-vaccinated group (**Fig 4f**). In contrast four out of six *ΔsigH* vaccinated macaques exhibited no signal in the thoracic cavity by PET/CT at the endpoint, and the other two macaques in this group exhibited minimal pathology, much of which was localized to the bronchial lymph node (**Fig 4g**). SUVmax values of *ΔsigH*-vaccinated group were significantly lower than both unvaccinated and BCG-vaccinated groups at this timepoint (**Fig 4h**). The number of lesions detected by PET/CT and volume occupied by them was also significantly lower at week 12-13 in both vaccination groups relative to unvaccinated macaques (**Fig S3c, d**). Therefore, the analysis of PET/CT scans clearly showed that *ΔsigH* vaccination protected macaques from *Mtb* Erdman challenge in an elite fashion relative to BCG.

**Figure 4.**
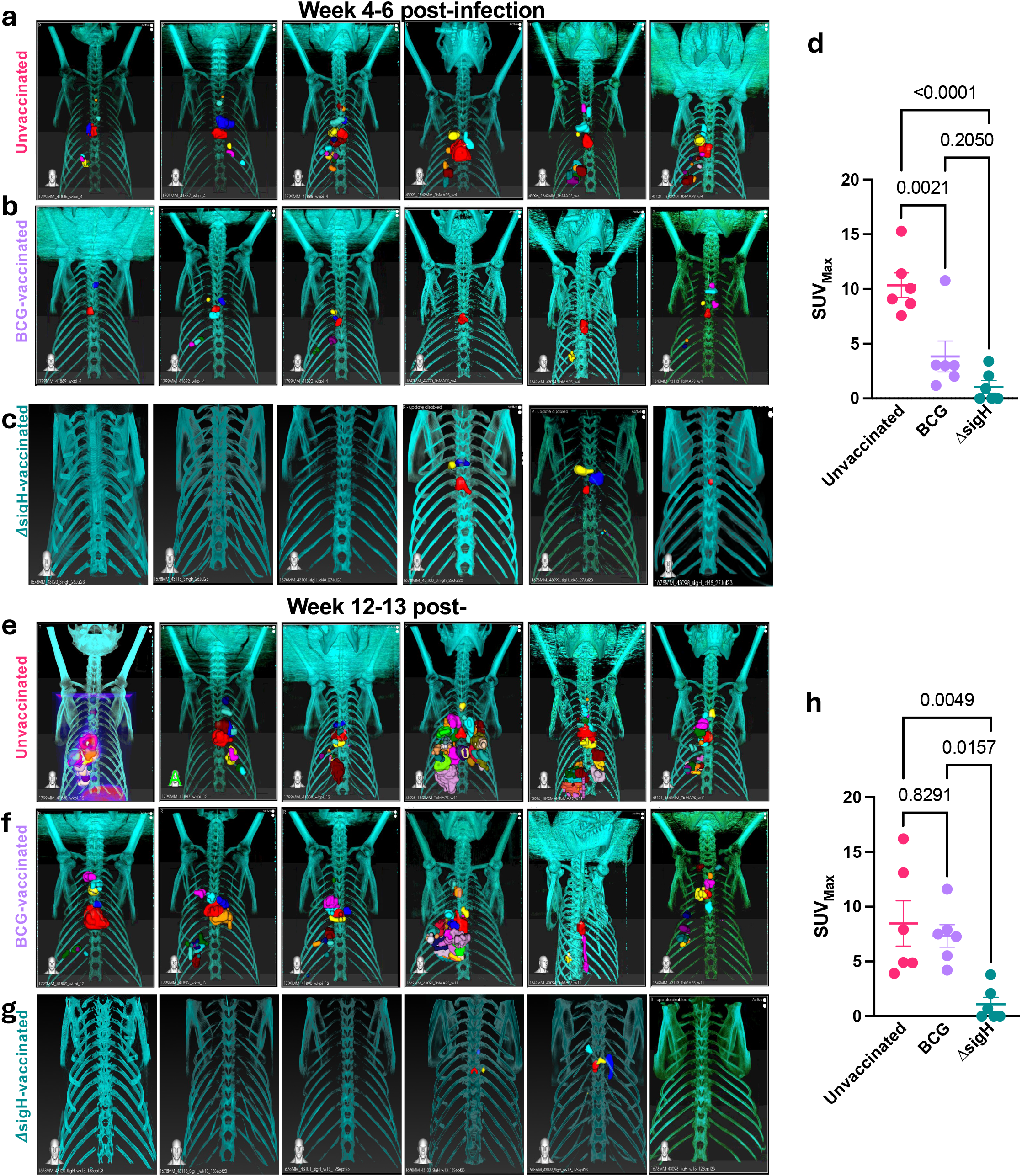

### *ΔsigH* vaccination induces elite antigen-specific T cell responses correlating with protection

To assess if the elite protection of susceptible rhesus macaques from virulent Mtb challenge by prior aerosol *ΔsigH* vaccination correlated with local antigen-specific T cell responses, cryopreserved BAL cells were stimulated with *Mtb* antigens and the expression of Th1, Tc1 and Th17 cytokines analyzed by flow cytometry (intracellular cytokine staining) and compared to responses elicited by BCG vaccination. Significantly greater frequencies of CD4^+^ T cells from the BAL of *ΔsigH* vaccinated macaques expressed IFNγ (**Fig 5a**), TNFα (**Fig 5b**), IL2 (**Fig 5c**), IL17 (**Fig 5d**) and both IFNγ and TNFα (**Fig 5f**) but not Granzyme B (**Fig 5e**), relative to BCG vaccinated macaques. Similarly, significantly greater frequencies of CD8^+^ T cells from the BAL of *ΔsigH* vaccinated, relative to BCG-vaccinated macaques, expressed IFNγ (**Fig 5g**), TNFα (**Fig 5h**), IL17 (**Fig 5j**) and both IFNγ and TNFα (**Fig 5l**). Although the frequencies of IL2 (Fig 5i) and Granzyme B (**Fig 5k**) expressing CD8+ T cells were also higher in the BAL of *ΔsigH*-vs BCG-vaccinated macaques, these differences were not statistically significant. Thus, as we have previously demonstrated in the resistant cynomolgus macaque model, *ΔsigH* vaccination also induces elite antigen-specific T cell responses correlating with protection in the susceptible rhesus macaque model.

**Figure 5.**
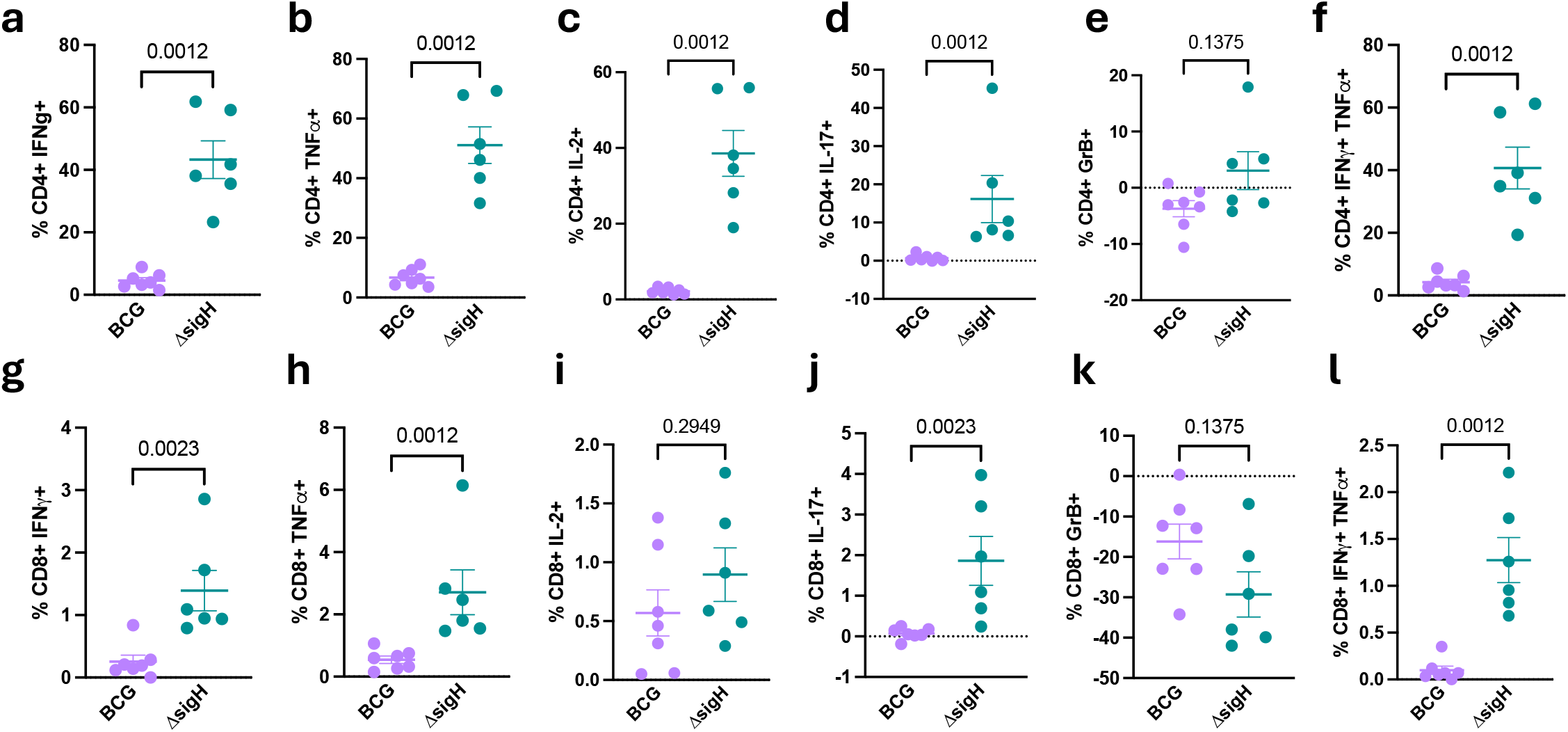

### Mucosal *ΔsigH* vaccination induces iBALTs, a tertiary lymphoid follicle associated with protective immunity

Inducible Bronchus-Associated Lymphoid Tissue (iBALT) consists of organized aggregates of B cells, T cells, dendritic cells, and macrophages that develop in the lung following antigenic insult usually due to an infection or vaccination. These ectopic follicular lymphoid tissues function similarly to secondary lymphoid organs by facilitating localized antigen presentation followed by lymphocyte activation and proliferation triggering rapid immune recall responses within pulmonary tissue. Our previous work in cynomolgus and rhesus macaques showed that aerosol vaccination with *ΔsigH* produced significantly greater iBALT formation compared with BCG. Histological analyses demonstrated large CD20^+^ B-cell follicles surrounded by CXCR5^+^ T cells within granulomatous lesions. These organized immune structures promoted enhanced recruitment and retention of lung-homing Th1/Th17 and Tc1/Th17 cells, resulting in improved macrophage activation and bacterial control. *ΔsigH*-vaccinated animals exhibited markedly reduced pulmonary pathology, lower bacterial burdens, and improved survival following virulent TB challenge ^23,24^. Single cell analysis of alveolar airway cells from *ΔsigH*-vaccinated macaques demonstrated higher complexity in T-B cell immune synapses compared to BCG^24^. To better understand the dynamics and functionality of these protective follicular iBALT structures, we performed 18F-FDG PET/CT imaging of all the *ΔsigH*-vaccinated macaques after six and eleven weeks of mucosal vaccination. PET/CT scans at week 6 post-vaccination demonstrated numerous follicular structures (**Fig 6a-c**) showing 18F-FDG uptake. These structures were metabolically active (**Fig 6a**), smaller in size than traditional granuloma (**Fig 6b**) and CT dense (**Fig 6c**). The iBALTs structures were observed throughout the study in PET/CT scans performed at week 11 post-vaccination (**Fig 6d-f**), Week 4-6 (**Fig 6g-i**) and 12-13 post-infection (**Fig 6j-l**). However, the number of iBALTs tends to increase until week 6 post-infection (**Fig 6m**) and decline towards the endpoint implying antigenic exposure as a potential factor required for maintenance and retention of these tertiary follicles. However, the observed metabolic activity in the iBALTs was highest during week 6 post-vaccination (**Fig 6n**) and transiently decreased over time. An example from one of the *ΔsigH*-vaccinated macaques is highlighted in **Fig 6o-p**. PET/CT scans performed between weeks 6-7 in these animals post-vaccination but before Mtb challenge show significant involvement of lung-draining lymph nodes but a complete lack of granulomatous pathology in the lung compartment (Fig 6o), thereby highlighting the attenuated but immunogenic nature of the *ΔsigH* strain. Instead, PET/CT identified these lungs to contain dozens of much smaller features, which corresponded to iBALT (**Fig 6p**). These data clearly demonstrate that iBALTs present as small hypermetabolic peribronchial lymphoid aggregates on PET-CT, their metabolic activity can be tracked over time non-invasively and usually correlates with the airway T cell antigenicity. These findings highlight that mucosal ***ΔsigH*** vaccination not only generates systemic immunity like traditional intradermal BCG vaccination but also actively remodel the lung immune microenvironment to create durable, tissue-resident protective immunity.

**Figure 6.**
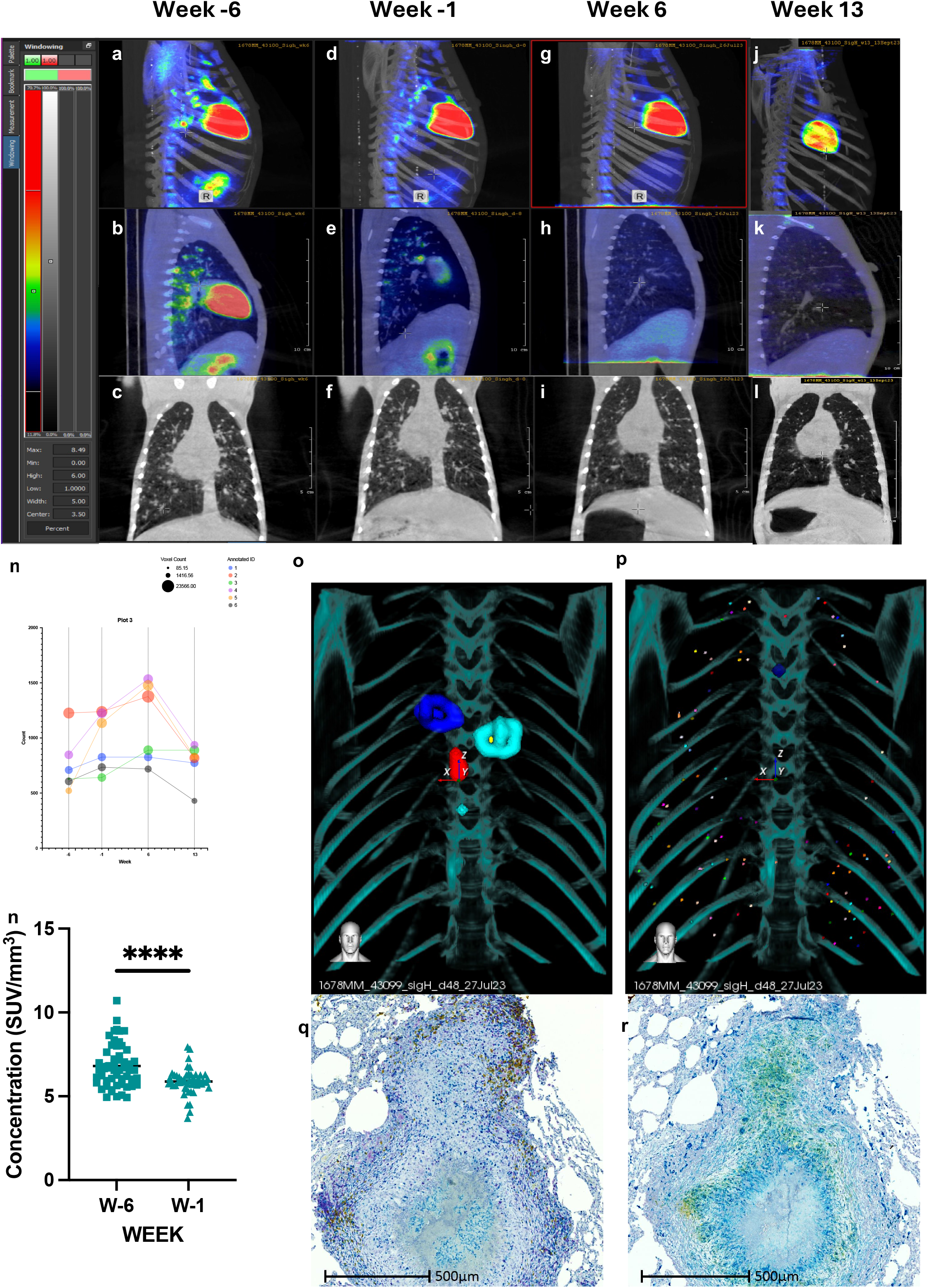

## Discussion

Live-attenuated *Mtb* vaccine candidates offer the advantage of presenting the immune system with a broad repertoire of TB antigens^37^. As a result, they can potentially induce comprehensive and more durable immune responses than vaccines that elicit responses to a limited number of antigens. Despite this, only one live-attenuated candidate - MTBVAC^38^ - has progressed to advanced clinical evaluation^39^. These observations highlight the need to develop additional rationally attenuated *Mtb* strains with improved immunogenicity and safety profiles. The *ΔsigH* mutant represents one such candidate. SigH is a master regulator of the mycobacterial stress response and protects bacilli from oxidative and other host-mediated stresses encountered during infection^9,10^. In the absence of SigH, *Mtb* is unable to neutralize host oxidative defenses, resulting in impaired survival within phagocytes and an inability to sustain bacillary replication in vivo. Rather than establishing productive infection, *ΔsigH* triggers robust innate and adaptive immune responses^18^, likely via enhanced antigen presentation by cells infected with *ΔsigH* compared with those infected with BCG or wild-type *Mtb* – a mechanism that we have experimentally demonstrated to be true^24^.

Our study demonstrates that aerosol vaccination with the live-attenuated *MtbΔsigH* mutant derived from the CDC1551 background confers protection against challenge with the heterologous and more virulent *Mtb* Erdman strain in the susceptible rhesus macaque model of TB. Vaccinated animals exhibited reduced pulmonary pathology, as measured by histopathology as well as PET/CT scanning, including fewer granulomatous lesions, and enhanced formation of lymphoid follicles consistent with inducible bronchus-associated lymphoid tissue (iBALT). Our earlier studies demonstrating *ΔsigH*-mediated protection used homologous challenge, where both the vaccine and the challenge strains were from the CDC1551 background. The present study addresses this limitation by demonstrating protection against heterologous challenge with the more virulent Erdman strain^25^. Thus, current findings demonstrate that *ΔsigH*-mediated immunity can protect against infection with a genetically and phenotypically distinct *Mtb* strain. This result is particularly relevant given the genetic diversity of circulating clinical *Mtb* strains worldwide. Despite the increased virulence of the Erdman strain and the heightened susceptibility of rhesus macaques relative to other NHPs, the extent of reduction in bacillary burdens observed in lung granulomas was comparable to that reported in our previous studies using CDC1551 challenge in either rhesus^23^ or cynomolgus^24^ macaques. These findings suggest that vaccination with *ΔsigH*-based strains may be capable of controlling *Mtb* replication even in settings where circulating strains exhibit increased virulence or drug resistance. These results strongly support the rationale for the further development of *ΔsigH*-based anti-TB vaccines, particularly since *ΔsigH* has previously been shown to be nonpathogenic in the setting of HIV-like immunosuppression using Simian Immunodeficiency Virus infection in rhesus macaques^19^. This observation raises the possibility that *ΔsigH*-based vaccines may ultimately be suitable for use in populations with high prevalence of HIV infection, who remain among the groups most vulnerable to TB^40^.

In addition to demonstrating that aerosol vaccination with *ΔsigH* protects against heterologous challenge, this study also significantly enhanced our mechanistic understanding of the immune mediators involved in engendering protection against lethal TB. We earlier showed that the lungs of rhesus macaques vaccinated with *ΔsigH* are characterized by the presence of greater iBALT at the post-*Mtb* challenge endpoint^23^. These results were confirmed in cynomolgus macaques vaccinated with *ΔsigH* by single cell multiplexed, spatial imaging^24^. scRNAseq evaluation of post-vaccination BAL of the same macaques showed signatures of enhanced T-B cell cooperation, potentially hinting at increased iBALT post-vaccination^24^. We also observed potent Type I IFN conditioning of T cells, but in the absence of pathological effects of Type I IFN signaling on myeloid cells^24^ that is associated with TB pathology^41^. Here, using whole-body imaging by PET/CT, we show for the first time, that high-dose aerosol *ΔsigH* vaccination induces iBALT responses post-vaccination, prior to *Mtb* challenge. Detected on PET/CT as small features distinct from granulomas, and in animals with potent lymph node metabolic activity, these iBALT are also highly metabolically active. Post-*Mtb* challenge, animals with iBALT do not develop granulomas or *Mtb* burdens in lungs, and metabolic activity of iBALT declines in this period. We have earlier shown that the depletion of B cells in *ΔsigH*-vaccinated rhesus macaques disrupts iBALT and significantly reduces the efficacy of vaccination^42^. Here, by demonstrating that the iBALT response is strongly generated by *ΔsigH* vaccination prior to *Mtb* challenge, we posit iBALT as the key mechanism by which mucosal vaccination may be used to elicit protective responses in individuals susceptible to lethal TB. Further characterization of iBALT via spatial imaging is currently ongoing.

Collectively, our findings provide further support for the continued development of *ΔsigH*-based live-attenuated *Mtb* vaccines. Demonstration of protection against heterologous challenge in a highly susceptible NHP model strengthens the translational potential of this approach and supports its continued evaluation in longer-term challenge studies and eventual clinical development. As specified by the Geneva consensus on the development of such vaccines^43,44^, further development of a *ΔsigH*-based anti-TB vaccine platform in the clinical realm would likely utilize multiple gene mutations in the *ΔsigH* background. We have developed eight such strains based on *ΔsigH*, seven of which are safe in both immunocompetent and immunodeficient macaques^45^.

Some limitations of our work remain. We have only tested the efficacy of *ΔsigH* to protect against lethal TB in a short-term challenge model. Clinical development of such candidates would require the demonstration of durable immunogenicity and efficacy in long-term challenge.

## Supporting information

All supplemental Figs

All supplemental Tables

## Author Contributions

DK designed the study; DKS, GA, VSRD, AD, MA, KW, VS, SH-U, ZL, SM and XA conducted research and/or performed data analysis; DK wrote the paper with help from DKS and SAK. DK and SAK provided funding. All authors contributed to and approved the manuscript.

## Acknowledgments

This research was supported by NIH grants AI111914, AI134240, AI138587 and AI184623 to DK and/or SAK and institutional NIH grants– OD010442, OD032443, OD028732, AI161943 and AI168439.

## Supplementary tables

**Supplementary Table 1: Demographics and legends for individual animals used in the study**.

**Supplementary Table 2: Flow Cytometry panels**

## Supplementary figures

**Fig S1. Post vaccination and post-challenge clinical, microbiological and pathology measures. a**. Antigen-specific intracellular cytokine staining pre-vaccination (turquoise) and post-challenge (orange) to measure IFNG production to confirm and validate Tuberculin Skin Test (TST) results, following stimulation with *Mtb* cell wall fraction (CW) (BEI Resources). **b**. % change in body weight at the end of the vaccination phase in BCG (lavender) and *ΔsigH*-vaccinated (teal) groups relative to pre-vaccination baseline. **c**. Peak serum C-reactive protein (CRP) values (mg/dL) during the vaccination phase in **BCG**- and ***Δ*sigH**-vaccinated groups. **d**. % change in serum A/G ratios at the end of the vaccination phase relative to pre-vaccination baseline in **BCG**- and ***Δ*sigH**-vaccinated groups. **e**. N/L ratio is peripheral blood at the end of the vaccination phase in **BCG**- and ***Δ*sigH**-vaccinated groups. **f**. CXR scores at the end of the vaccination phase in **BCG**- and ***Δ*sigH**-vaccinated groups. **g**. BAL CFUs at the end of the vaccination phase in **BCG**- and ***ΔsigH****-*vaccinated groups. **h** (peak) and **i** (endpoint) serum CRP values (mg/dL) in unvaccinated (strawberry), BCG (lavender) and *ΔsigH*-vaccinated (teal) groups during the *Mtb* challenge phase. Changes in blood neutrophil (N) percentage (**j**) and numbers (**k**) in **unvaccinated, BCG**- and ***ΔsigH***-vaccinated groups at necropsy. (**l**). BAL CFUs (at the endpoint) in **unvaccinated, BCG**- and ***ΔsigH***-vaccinated groups. CFUs in spleen (**m**), liver (**n**) and kidneys (**o**) (per gram tissue) in **unvaccinated, BCG**- and ***ΔsigH***-vaccinated groups at necropsy. **(q)** Fisher’s exact test comparing sterile (strawberry) vs nonsterile (turquoise) lung lobes in the three groups of macaques. Data is presented as mean ± SEM and P-values are derived from multiple Mann-Whitney tests with multiple hypothesis correction by false discovery rate method of Benjamini, Kreiger and Yekutielli (two stage step up) (**b-g**) except for **h-o** where one-way ANOVA with Tukey’s correction was used.

**Fig S2. Lymphocytic immune responses in BAL post-infection**. In three groups of CMs, shown are the absolute counts of CD3^+^ (**a**), CD4^+^ (**b**), and CD8^+^ (**c**) T cells and B cells (**d**), in BAL at weeks 3, 5, and 7 post-infection time-point. Each column represents an individual macaque. Results are shown for n=5 macaques per group. Shown are the frequencies of CD3^+^CCR5^+^ (**e**), CD4^+^CCR5^+^ (**f**), and CD8^+^CCR5^+^ (**g**) T cells in BAL, expressed as percentage of parental populations. Frequencies of CD4^+^ T cells (CD28^+^CD95^-^) effector (**h**), (CD28^+^CD95^-^) memory (**i**) and naïve (CD28^-^CD95^-^) (**j**), CD8^+^ T cells (CD28^+^CD95^-^) effector (**k**), (CD28^+^CD95^-^) memory (**l**) and naïve (CD28^-^CD95^-^) (**m**) in BAL at weeks 3, 5, and 7 post-infection time-point, expressed as percentage of parental population. All frequency results are shown for weeks 3, 5, and 7 post-vaccination time-point. Shown are the frequencies of CD20^+^CCR5^+^ (**n**), CD20^+^CCR7^+^ (**o**), CD20^+^KI67^+^ (**p**) and CD20^+^CD69^+^ (**q**) B cells in BAL, expressed as percentage of B cell population. Each column represents an individual macaque. Results are shown for n=5 macaques per group.

## Notes

<u>Conflict of interest’s statement:</u> “The authors have declared that no conflict of interest exists.”

### Competing Interest Statement

The authors have declared no competing interest.

