## Supplementary figures and images for "Bronchus associated lymphoid tissue induced by an attenuated *Mycobacterium tuberculosis* vaccine prevents tuberculosis from heterologous TB challenge"

### All supplemental Figs

#### Slide 1
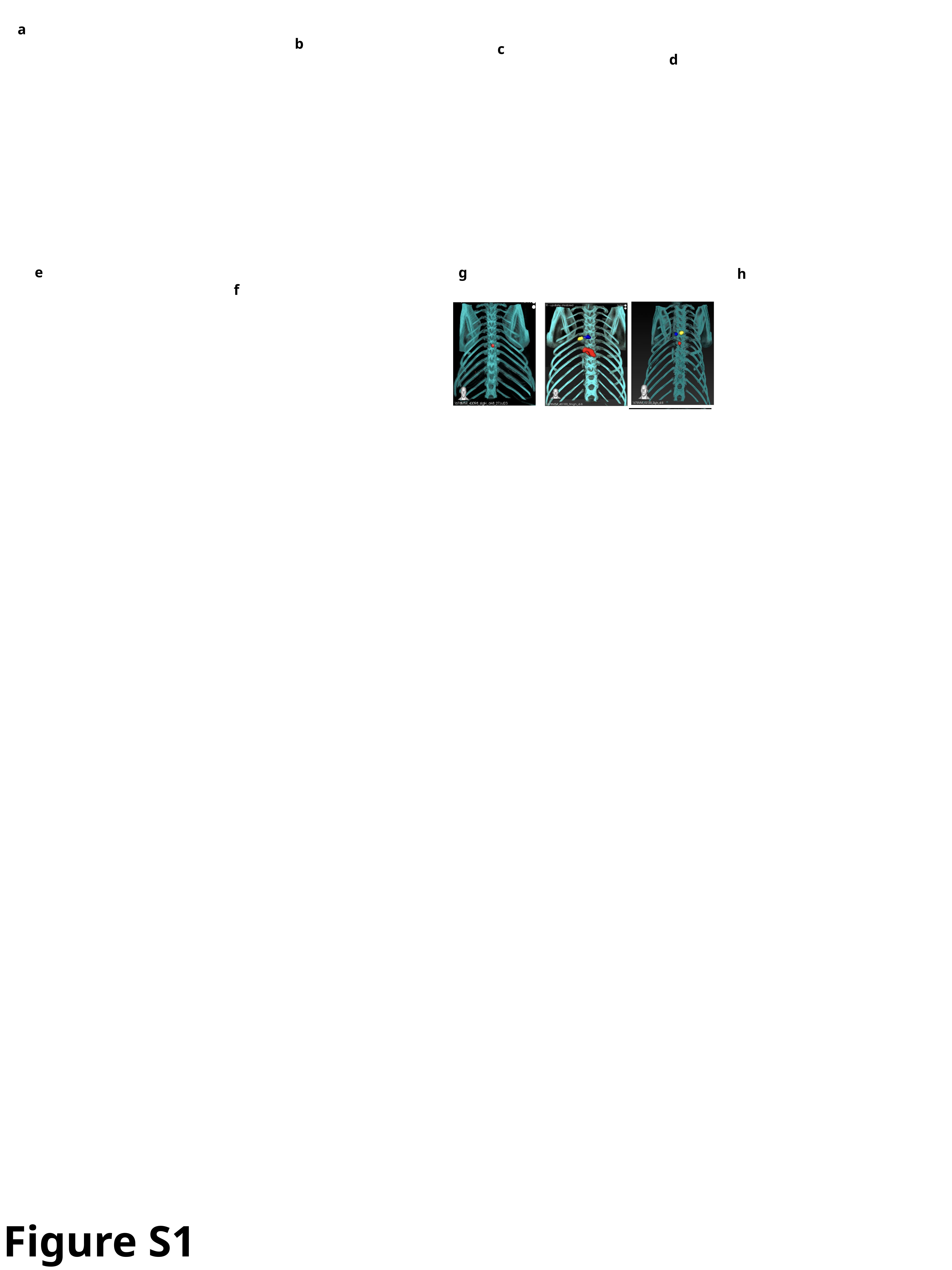

a
b
c
d
e
g
f
h
Figure S1

#### Slide 2
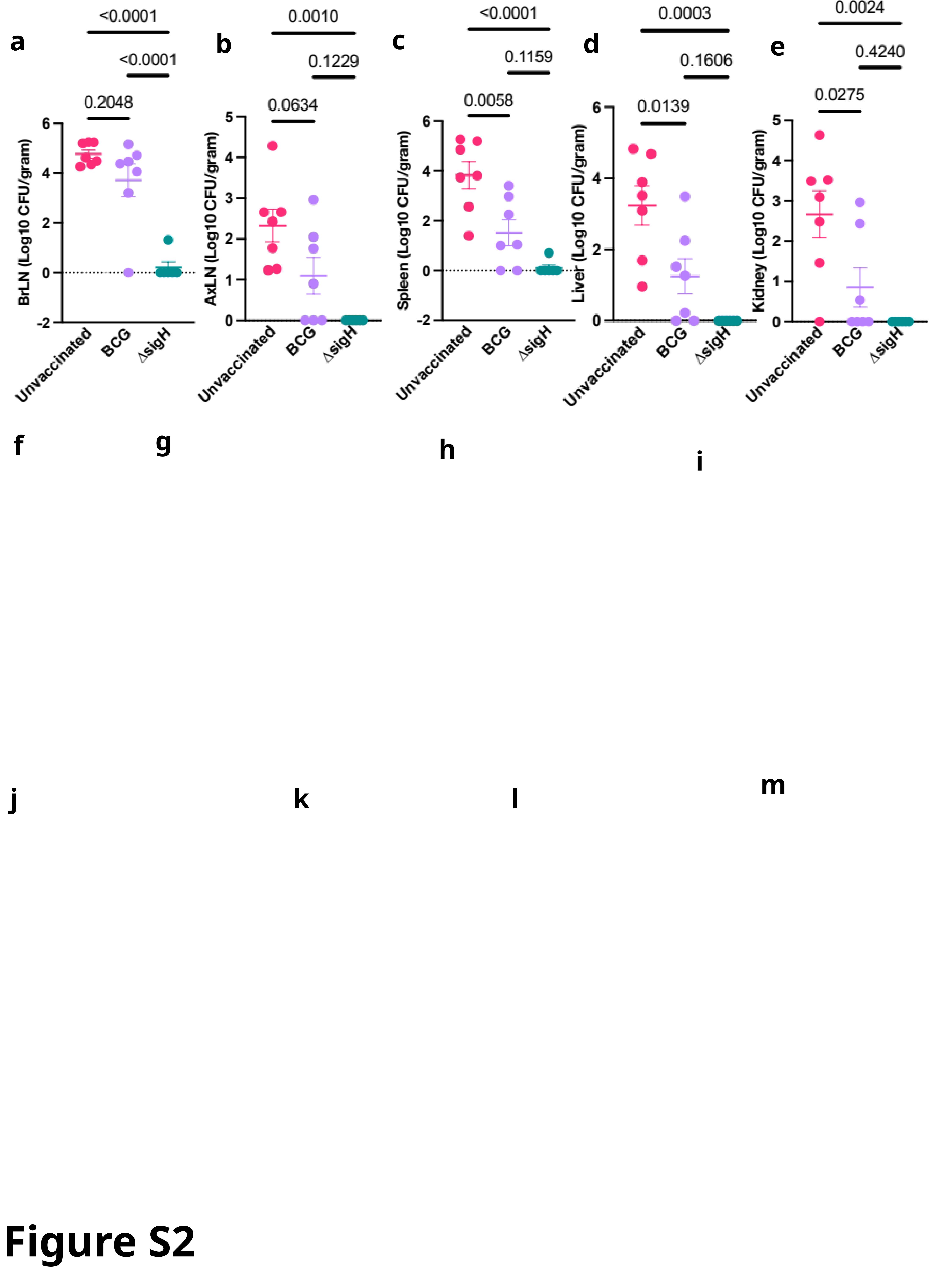

c
a
d
b
e
g
f
h
i
m
j
k
l
Figure S2

#### Slide 3
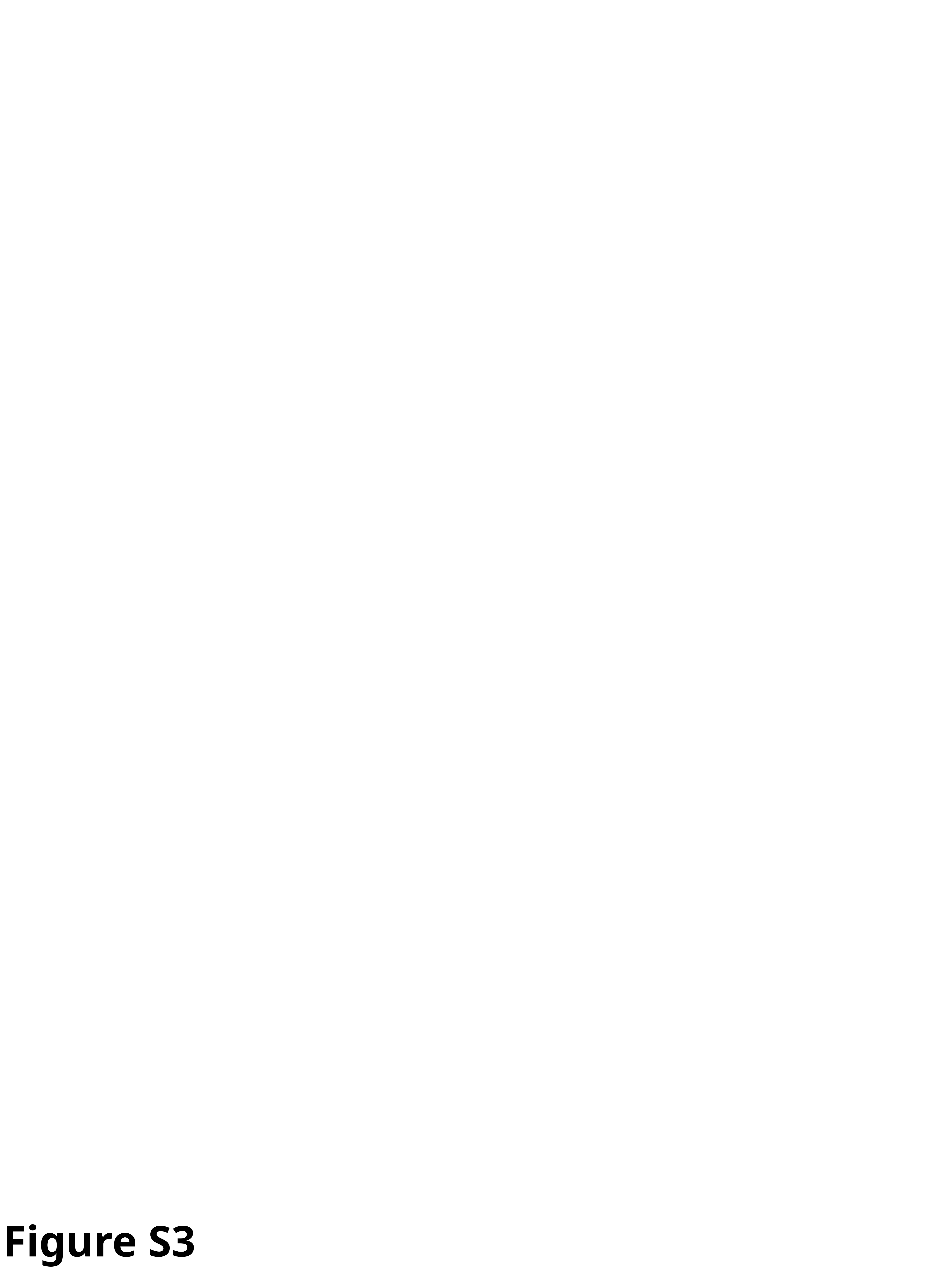

Figure S3
